# Human malaria parasite cold shock protein plays an essential role in asexual and sexual stage development and presents an excellent druggable target

**DOI:** 10.1101/2022.07.07.499247

**Authors:** Ankita Behl, Monika Saini, Fernando De Leon, Geeta Kumari, Vikash Kumar, Rumaisha Shoaib, Harshita Singh, Swati Garg, Christoph Arenz, Shailja Singh

## Abstract

Cold shock proteins are well characterized in bacteria, plants and humans, however there is no information on their existence and role in malaria parasite. Here, we have delineated the function of a novel cold shock protein of *Plasmodium falciparum* (*Pf*) which we have annotated ‘*Pf*CoSP’. Our results show that recombinant *Pf*CoSP has both DNA and RNA binding activity, and also interacts with alpha and beta tubulin of *Pf. Pf*CoSP binds with RNA and alpha tubulin simultaneously to form a complex. Expression of *Pf*CoSP was found during asexual blood stages and gametocyte stages of malaria parasite. *Pf*CoSP expression up-regulates many folds upon cold treatment, suggesting its role during hypothermic cold shock. Interestingly, *Pf*CoSP showed binding with human cold shock protein LIN28A inhibitor ‘LI71’ that also inhibits *Pf*CSP -alpha/beta tubulin interactions. LI71 showed antimalarial activity against asexual blood stage and gametocyte stage suggesting its multi-stage, transmission-blocking potential. We propose that *Pf*CoSP may form a workbench for translation during cold stress via its interactions with target mRNAs and the cytoskeleton protein tubulin.

## Introduction

All organisms face changing environmental conditions and have the potential to cope with different types of stress environments. Cold shock is one of the most common stresses faced by all living organisms. To cope with adverse effects of low temperature, cold shock proteins are expressed in bacteria, plants and humans that play a significant role in acquiring cold tolerance [1-5]. Cold shock proteins harbor one or more cold shock domain that forms the hallmark of cold shock protein family. Cold shock domain has DNA and RNA binding motifs that make cold shock proteins regulate critical processes inside the cell including transcription, translation and splicing. Cold shock proteins are known to destabilize secondary structures in target RNA which in turn allow efficient transcription and translation [3, 6].

Cold shock proteins were initially identified in bacteria when a drop in temperature caused many fold increase in expression of a cold shock protein A (CspA) in *Escherichia coli* [7, 8]. *Escherichia coli* encodes 9 cold shock protein genes (*CspA* to C*spI*) that encode highly conserved family of structurally related proteins with a molecular mass of approximately 7.4 kDa [9]. Bacterial Csps have typical cold shock domain that bestow them with the ability to bind with single-stranded RNA and DNA [10-14]. This protein-DNA/RNA binding is mediated by moderately well-conserved nucleic acid binding motifs RNP1 (K/R-G-F/Y-G/A-F-V/I-X-F/Y) and RNP2 (L/I-F/Y-V/I-G/K-N/G-L) [15-16]. Structural studies on bacterial cold shock proteins indicate that they possess an OB (oligonucleotide/oligosaccharide-binding) fold that comprises 5 antiparallel beta strands forming a Greek-key beta-barrel [17].

In humans, eight members of cold shock proteins exist that are encoded by genes namely YBX1, YBX2, YBX3, CARHSP1, CSDC2, CSDE1, LIN28A, and LIN28B [18]. YBX1, YBX2, YBX3 encode Y-box binding protein-1 (YB-1), DNA binding protein A (DbpA) and C (DbpC) respectively and comprise Y-box binding protein family [18]. YB-1 binds with both DNA and RNA and is involved in several critical processes including mRNA splicing, mRNA translation, DNA replication and repair [19-22]. Calcium-regulated heat-stable protein 1 (CARHSP1) is known to bind with tumor necrosis factor (TNF) mRNA [23] whereas PIPPi (encoded by CSDC2) binds specifically to the 3’-UTR ends of both histone H1 and H3.3 mRNAs [24]. Role of CARHSP1 is associated with TNF stabilization within P-bodies and exosomes [23] while PIPPin is implicated in negative regulation of histone variant synthesis in the developing brain [24]. Unr (upstream of N-ras) is another human cold shock protein (encoded by CSDE1) that is involved in translational reprogramming [25]. Another developmentally important cold shock protein expressed in humans is Lin28 whose function is to regulate translation of mRNAs that control developmental timing, pluripotency and metabolism [26].

Cold shock proteins are also well characterized in plants like wheat, rice and *Arabidopsis thaliana* and are known to perform several functions like regulating flowering time, embryo and fruit development and also help plants in acquiring freezing tolerance [27]. Radkova *et al*. identified 4 cold shock protein genes in wheat that were further classified into three classes [28]. Expression of class I and class II proteins is upregulated during flower and seed development while class III wheat cold shock protein (27 kDa) is expressed only during seed development [28]. *Arabidopsis thaliana* has 4 cold shock proteins 1-4 (AtCSP1-CSP4) that show binding with RNA, single and double-stranded DNA. Among these, AtCSP3 (At2 g17870) is essential for acquiring freezing tolerance [29] whereas AtCSP2 and AtCSP4 are involved in regulating development processes [30, 31]. Chaikam and Karlson identified two cold shock proteins in rice (OsCSP1 and OsCSP2) whose role is more related with developmental processes as compared to cold tolerance [32].

Although the functions of cold shock proteins are well known in bacteria, plants and humans, there is no information on the presence and role of cold shock proteins in *Plasmodium falciparum*. Our search in Plasmodb database [33] identified a single cold shock gene in malaria parasite. The PhenoPlasm database [34] suggested that *Pf*CoSP is essential for survival of the parasite inside the human host. This underscores its importance in parasite biology and makes it an attractive candidate for development of anti-malarials. In the present study, we have attempted to elucidate the function of *Pf*CoSP in malaria parasite. Our data suggest that *Pf*CoSP has nucleic acid binding ability and interacts with alpha and beta tubulin. *Pf*CoSP forms a complex by binding with RNA and alpha tubulin simultaneously. The expression of *Pf*CoSP was found mainly in trophozoite and schizont stages and upregulates upon cold stress in malaria parasite. Also, *Pf*CoSP binds with LIN28 inhibitor ‘LI71’ that inhibits *Pf*CSP -alpha/beta tubulin interactions and showed antimalarial activity against *Pf*3D7 and *Pf*RKL-9. Overall, these data indicate that *Pf*CoSP may play a role in growth and development of malaria parasite upon cold stress.

## Material and methods

### Parasite culture

*Plasmodium falciparum* 3D7 parasites were cultured in O+ RBCs using complete RPMI 1640 medium supplemented with 0.5 gm/L Albumax I (Gibco, USA), 27.2 mg/L hypoxanthine (Sigma, USA) and 2 gm/L sodium bicarbonate (Sigma, USA). Culture was maintained at 37°C in 90% N_2_, 5% CO_2_, and 5% O_2_ containing environment and maintained at 5% haematocrit and 5% parasitaemia. Cultures were harvested by centrifugation from *Plasmodium* cultures (Parasetemia 8-10%) and the parasites were released from red blood cells by treatment with 0.15% saponin. Parasite pellet was washed with 1X PBS and stored at -80 °C for experiments. RNA was isolated from infected RBCs using TRIzol reagent. DNA was removed by the DNase treatment kit (DNA-*free*™ DNA Removal Kit, Invitrogen, Thermo Fisher Scientific, Massachusetts, USA). Parasites were synchronized by 5% sorbitol (Sigma) selection of rings and late trophozoites or schizonts were purified from mixed parasite culture using 65% Percoll (Sigma).

### Molecular cloning, over-expression and purification of *Pf*CoSP

Full-length gene of PF3D7_0109600 (*Pf*CoSP) was amplified by PCR using specific primers and *P. falciparum*-cDNA as template before cloning into bacterial expression vector pET28a. After screening the transformed *E. coli* DH5α cells for positive clones by restriction digestion, recombinant vector carrying sequence for *Pf*CoSP was transformed in BL21 (DE3) *E. coli* cells. Expression of *Pf*CoSP was induced with 1 mM IPTG at 37ºC for 4 hrs. Purification of recombinant protein was achieved by affinity chromatography.

### Raising polyclonal antisera against *Pf*CoSP

Polyclonal antibodies were raised in house against *Pf*CoSP in male BALB/c mice using purified recombinant protein as immunogens following standard protocol [35]. Briefly, animals were immunized with emulsion containing 1:1 ratio of freund’s complete adjuvant and antigen followed by three booster doses at an interval of 14 days. Booster doses comprise 1:1 ratio of freund’s incomplete adjuvant and antigen. Final bleed was collected and the end point titers of raised antisera were determined by western blotting.

### Estimation of number of transformed *E. coli* cells upon temperature stress

BL21 (DE3) *E. coli* cells transformed with cloned plasmid of *Pf*CoSP was induced with 1mM IPTG for 4 hrs and then subjected to temperature stress of 25°C for 8 hrs. Cultures were taken at 4 hrs and 8 hrs post cold treatment and subjected to kanamycin plates at different dilutions. Plates were kept at 37°C overnight.

### Gel retardation assay

DNA/RNA binding property of protein was determined by gel retardation assay. Purified recombinant protein (1μgm) was incubated with 100ng *Pf*DNA and *Pf*RNA in 2X nucleic acid binding buffer (3 mM MgCl_2,_ 50% glycerol, 1M HEPES, 100 mM DTT, 1M tris-Cl pH 8) at 37°C for 1 hr. Migration patterns of nucleic acid samples incubated with protein was studied by agarose gels stained with ethidium bromide.

### Bead based *Pf*CoSP-DNA pull down assay

The nucleic acid binding property of protein was further validated using cellulose beads with immobilized single and double stranded calf thymus DNA (Sigma Aldrich, USA). 1 μgm recombinant protein was mixed with 500 µl cellulose matrix (1mg/ml) and incubated at 4°C for 30 mints. Bead pellet was washed with 200 µl buffer and were boiled in 1X SDS-PAGE sample loading dye before loading on 15% SDS-PAGE.

### Binding and inhibition assays using Nanotemper

The kinetic measurements of *Pf*CoSP-gDNA binding were conducted using Monolith NT.115 instrument (NanoTemper Technologies, Munich, Germany). 10 μM *Pf*CoSP was labeled using NanoTemper’s Protein Labeling Kit RED-NHS (L001, NanoTemper technologies, Germany). The concentration of labeled *Pf*CoSP protein was kept constant at a concentration of 90 nM. The unlabeled binding partners (genomic DNA of *Pf*) were titrated in 1:1 dilution. 2pM *Pf*gDNA and 100nM *Pf*RNA was serially diluted with decreasing concentrations and was titrated against constant concentration of the labelled *Pf*CoSP. Samples were diluted in 1x PBS buffer. For measurements, samples were filled into standard treated capillaries.

We also evaluated interaction of α-tubulin and β-tubulin with *Pf*CoSP using Monolith NT.115 instrument. 20 µM each of α-tubulin and β-tubulin recombinant proteins in 1X PBS buffer, pH 7.5, were fluorescently labelled using 30 μM Lysine reactive dye (Monolith ™ Series Protein Labelling Kit Red-NHS 2^nd^ Generation), followed by incubation for 30 minutes in dark at room temperature. The labelled proteins were passed through an equilibrated column (provided in the kit) and elution fractions were collected followed by subjecting the elutions to fluorescence count. Simultaneously, 30 µM recombinant *Pf*CoSP was serially diluted with decreasing concentrations in 1X PBS/0.01% Tween20 and was titrated against constant concentration of the labelled α-tubulin and β-tubulin proteins. Samples were incubated for 15 minutes at room temperature, followed by centrifugation at 8000 rpm for 10 minutes at room temperature. The samples were taken into the capillaries (K002 Monolith NT.115) and thermophoretic mobility was analyzed.

Similar protocol was followed to investigate the interaction of *Pf*CoSP with LI71. 10 μM *Pf*CoSP was labeled and the concentration of labeled *Pf*CoSP was kept constant at a concentration of 626nM. The unlabeled binding partner LI71 was titrated in 1:1 dilution. The concentration of LI71 was 0.00061 µM, 0.00122 µM, 0.00244 µM, 0.00488 µM, 0.00977 µM, 0.0195 µM, 0.0391 µM, 0.0781 µM, 0.156 µM, 0.0391 µM, 0. 0781 µM, 0.156 µM, 0.313 µM, 0.625 µM, 1.25 µM, 2.5 µM, 5 µM, 10 µM, 20 µM. Samples were diluted in 0.01% Tween 20/1x PBS buffer.

We further performed competition experiments using MST to analyse affinity-based interaction of ligand molecules α-tubulin and β-tubulin competing with LI71 to interact with *Pf*CoSP. To serve this purpose, labelled α-tubulin and β-tubulin were mixed with 10 µM *Pf*CoSP at room temperature and incubated for 15 minutes. To this stock solution, serially diluted compound LI71 starting at 20 µM was added in equal amount. The samples were processed further as mentioned above. Data evaluation was performed with the Monolith software (Nano Temper, Munich, Germany).

### Plate based interaction studies

*In vitro* interactions of *Pf*CoSP with alpha and beta tubulin were examined by indirect ELISA. 100 ng each of purified alpha and beta tubulin were coated on ELISA plates and blocked with 5% BSA in PBS overnight at 4°C. The coated ligands were incubated with increasing concentrations of *Pf*CoSP (range: 10 ng to 500 ng) for 2 hrs at room temperature, followed by extensive washing with 1X PBS. Washed plates were incubated with anti-*Pf*CoSP (1:5000) followed by anti-mice HRP conjugated secondary antibodies (1:10,000) for two hrs. Plates were developed using 1mg/ml OPD (o-phenylenediamine dihydrochloride) containing H_2_O_2_, and absorbance measured at 490 nm.

### Co-immunoprecipitation assay

Pierce chemical co-immunoprecipitation kit was used according to the manufacturer’s protocol to evaluate native interaction of *Pf*CoSP and alpha and beta tubulin. Briefly, 10 µl of coupling resin was cross-linked with anti-*Pf*CoSP and anti-alpha/ beta tubulin antibodies for 2 hrs. After extensive washing, antibody coupled resin was incubated with parasite lysate. Unbound proteins were removed by repeated washing with binding buffer, followed by elution of prey proteins. As a negative control, preimmune sera were coupled to the resin before incubation with parasite lysate. Samples of bound proteins were resolved on 12% SDS-PAGE and subjected to western blot analysis using anti-*Pf*CoSP and anti-alpha/ beta tubulin antibodies.

### Complex binding assay

*In-vitro* formation of a complex between *Pf*CoSP, RNA/DNA and alpha/beta tubulin were tested using cellulose beads with immobilized single and double stranded calf thymus DNA (Sigma Aldrich, USA). 1 µg of *Pf*CoSP was mixed with 500 µl cellulose matrix (1mg/ml) and incubated at 4°C for 30 mints. Bead pellet was extensively washed with 200 µl buffer followed by incubation with 5 µg recombinant purified alpha/beta tubulin. After washing, beads were boiled in 1X SDS-PAGE sample loading dye before loading on 12% SDS-PAGE. Immunoblotting was performed to probe for *Pf*CoSP and alpha/beta tubulin using anti-*Pf*CoSP and anti-alpha/ beta tubulin antibodies respectively on the same blot. *Pf*CoSP was incubated with cellulose beads with immobilized single and double stranded calf thymus DNA separately as positive controls. BSA was incubated with beads separately and incubated with alpha/beta tubulin as negative controls.

### Real time PCR

Expression of *Pf*CoSP was evaluated at transcript level in asexual blood stages of *Pf*3D7 by real time PCR (StepOnePlus Real time PCR system Applied Biosystems, USA). The 18S rRNA gene of *Plasmodium falciparum* was chosen as a positive control. 10-μl reaction mixture comprises 1 μl of cDNA (1 μg), 5 μl of SYBR™ Green PCR Master Mix (Applied Biosystems™) and 1 μl (5 mM) of *Pf*CoSP specific forward and reverse primer. The PCR conditions consisted of an initial denaturation at 95°C for 5 min, followed by amplification for 40 cycles of 15 sec at 95°C, 5 sec at 55°C, and 1 min at 72°C, with fluorescence acquisition at the end of each extension step. Amplification was immediately followed by a melt program consisting of 15 sec at 95°C, 1 min at 60°C, and a stepwise temperature increase of 0.3°C/s until 95°C, with fluorescence acquisition at each temperature transition.

### *In-vivo* expression analysis of *Pf*CoSP

For *in vivo* expression analysis, *Pf*3D7 mixed stage asexual cultures (parasitemia ∼8%) were subjected to saponin lysis (0.15% w/v) followed by extensive washing of parasite pellet with 1x PBS to remove traces of haemoglobin. Protein extract of infected and uninfected RBCs (negative control) were resolved on SDS-PAGE and transfer to NC membrane followed by blocking in 5 % BSA for overnight at 4ºC. Following blocking, the membrane was washed with PBS (Phosphate buffer saline) followed by probing with specific antisera against *Pf*CoSP (1:4000) for 2 hrs. Post washing with PBST (0.025%) and PBS, NC membrane was hybridized with horseradish peroxidise (HRP)-conjugated secondary antibodies for 2 hrs. Membrane was subsequently washed with PBST (0.025%) and PBS, and developed with DAB/H_2_O_2_ substrate.

### Cold shock assay

Synchronised schizont-stage cultures at 4% haematocrit and 8% parasitaemia were exposed to temperatures at 25°C, 37°C and 42°C for 6 hours. Parasite pellet was boiled in 1X SDS-PAGE sample loading dye before loading on 15% SDS-PAGE. Samples were subjected to western blot and probed with anti-CSP antibodies. Blots were developed with DAB/H_2_O_2_ substrate.

### Immunofluroscence assays

Thin blood smears of mixed stage *Pf*3D7 cultures at 5% parasitaemia were fixed in methanol for 45 min at -20 °C, permeablized with 0.05% PBS/Tween 20, and blocked with 5% (w/v) BSA in PBS. For co-localization studies, mouse anti-*Pf*CoSP (1:250) and rabbit anti-PfNapL (1:250) and rabbit anti-tubulin (1:200) were added as primary antibodies and incubated for 2 hrs at room temperature. Alexa Fluor 488 conjugated anti-rabbit (1:500, red colour, Molecular Probes, Invitrogen, Carlsbad, CA, USA) and Alexa Fluor 546 conjugated anti-mouse (1:500, green colour; Molecular Probes) were used as secondary antibodies. The parasite nuclei were counterstained with DAPI (40, 60-diamidino-2-phenylindole; Invitrogen, USA) and mounted with a coverslip. The slides were examined using a confocal microscope (Olympus, Shinjuku, Tokyo, Japan) with a 9 100 oil immersion objective.

### *In vitro* antimalarial activity of LI71 against human malaria parasite

The antimalarial activity of LI71 was tested on *P. falciparum* 3D7 and RKL-9. Different concentration (250, 125, 62.5, 31.25, 15.62, 7.8, 3.9, 1.95 nM) of compound were added in 96-wells flat-bottom microplates in duplicate. Sorbitol synchronized cultures with 0.8-1% parasitemia and 2% hematocrit were dispensed into the plates and incubated for 72 h in a final volume of 100 µL/well. Chloroquine was used as a reference drug. Parasite growth was determined with SYBR Green I based fluorescence assay. Briefly, after 72 h of incubation culture was lysed by freeze-thaw followed by addition of 100 µL of lysis buffer (20□M Tris/HCl (Sigma-Aldrich, St. Louis, MO, United States), 5□M EDTA (Sigma-Aldrich, St. Louis, MO, United States), 0.16% (w/v) saponin (Sigma-Aldrich, St. Louis, MO, United States), 1.6% (v/v) Triton X (Sigma-Aldrich, St. Louis, MO, United States)) containing 1× SYBR Green I ((Thermo Fisher Scientific, Waltham, Massachusetts, US)). Plates were incubated in the dark at room temperature (RT) for 3-4□h. *P. falciparum* proliferation was assessed by measuring the fluorescence using a Varioskan™ LUX multimode microplate reader (Thermo Scientific™) with an excitation and emission of 485□nm & 530□nm respectively. IC_50_ values were determined via non-linear regression analysis using GraphPad prism 8.0 software. The results were expressed as the percent inhibition compared to the untreated controls and calculated with the following formula: 100× ([OD of Untreated sample – blank]-[OD– blank]/ [OD – blank]). As blank, uninfected RBCs were used. IC_50_, which is the dose required to cause 50% inhibition of parasite viability, was determined by extrapolation.

### Cold shock *P. falciparum* growth assays

Synchronous cultures of *P. falciparum* (3D7) were diluted to 1% parasitaemia and dispensed in 96-well plates along with different concentration of LI71 ((250, 125, 62.5, 31.25, 15.62, 7.8, 3.9, 1.95 nM). For cold shock treatment plates were incubated at 21°C for 4 h. Media was exchanged post treatment shock and incubated for 72 h. For control parasites were treated for 4 h with compound at different concentration and incubated at 37°C followed by washing and media exchanged (no drug). SYBR Green-I based fluorescence assay was performed to quantify the parasitemia. IC_50_ values were determined from growth inhibition data using nonlinear regression analysis (Prism 8, GraphPad). All data represent means of results from 3 independent experiments using biological replicates.

### *P. falciparum* gametocyte culture and gametocytogenesis

The strains of RKL9 of *P. falciparum* were cultured *in vitro* as described by Trager and Jensen with minor modifications. Briefly, parasites were maintained in human type 0 positive RBCs at 5% haematocrit (HCT) in RPMI 1640 medium containing 2 g/L sodium bicarbonate, 50 mg/L hypoxanthine (Sigma-Aldrich, St. Louis, MO, United States) with the addition of 10% (v/v) naturally clotted heat-inactivated 0+ human serum. The cultures were maintained at 37°C in a standard gas mixture consisting of (5% O2, 5% CO2, and 90% N2). To trigger gametocytogenesis, 10 mL cultures were diluted to 0.5% parasitaemia at 5% HCT. Media was changed daily for 3 days until 5% parasitaemia was reached and unhealthy, starved asexual parasites were detectable. At this point the hct was lowered to 2.5% and the culture was treated for 48–72 h with 50 mM N-acetylglucosamine (NAG; Sigma-Aldrich) in order to clear residual asexual parasites and obtain a virtually pure gametocyte culture, typically containing 2–4 gametocytes/100 RBCs at late stage II to early stage III of maturation.

### Gametocyte maturation in microwell plates

To analyse gametocyte development in 96-well plates, asexual parasite cultures were synchronized by purifying schizonts percoll gradient-centrifugation and allowed to reinvade erythrocytes. After two rounds of reinvasion, the cultures were treated with NAG (50 mM) for 72 h in order to clear residual asexual parasites and obtain a virtually pure gametocyte culture. Aliquots of 100 μL of synchronized gametocyte culture (typically at ∼2%gametocytaemia), diluted to 1% HCT, were seeded in 96-well flat-bottomed plates in presence of LI71 at 250 nM and 500 nM concentration respectively. Gametocyte were left untreated as control. Media with appropriate concentration of LI71 was exchanged daily for continuous 12 days. Giemsa-stained blood smears were observed to see the effect of LI71 on gametocyte development.

### Effect of L171 on *P. falciparum* exflagellation *in vitro*

To study the effect of LI71 on *P. falciparum* male gametocyte ex-flagellation, stage V gametocytes were treated for 30 min with 250 and 500 nM of LI71. Infected RBCs were left untreated as control. Samples were kept at 37ºC and incubated for 1 h. After incubation compound treated samples were washed and mixed immediately with 200 μL of ex-flagellation medium (RPMI1640 containing 25 mM HEPES, 20% FBS, 10 mM sodium bicarbonate and 50 mM xanthurenic acid at pH 7.4) and kept at 24ºC for 15 min. Ex-flagellation centres were then counted in 10-12 field using 40X objective of Fluorescence microscope.

## Results

### Domain organization, cloning, expression and purification of *Pf*CoSP

Sequence analysis and domain organisation using Blastp suggested that *Pf*CoSP is 150 amino acid long and harbor a typical N-terminal CSD domain containing DNA binding site and RNA binding motif (Fig. 1A). *Pf*CoSP was cloned in T7 promoter based plasmid pET-28a(+) and expressed in the soluble form in *E. coli* BL21 (DE3) cells with a 6X hexahistidine tag. *Pf*CoSP was purified using Ni-NTA affinity chromatography (Fig. 1B, left panel). Identity of recombinant *Pf*CoSP was checked by western blotting using anti-histidine antibodies (Fig. 1B, right panel). Polyclonal antisera were raised in house against *Pf*CoSP in male BALB/c mice.

**Figure 1:**
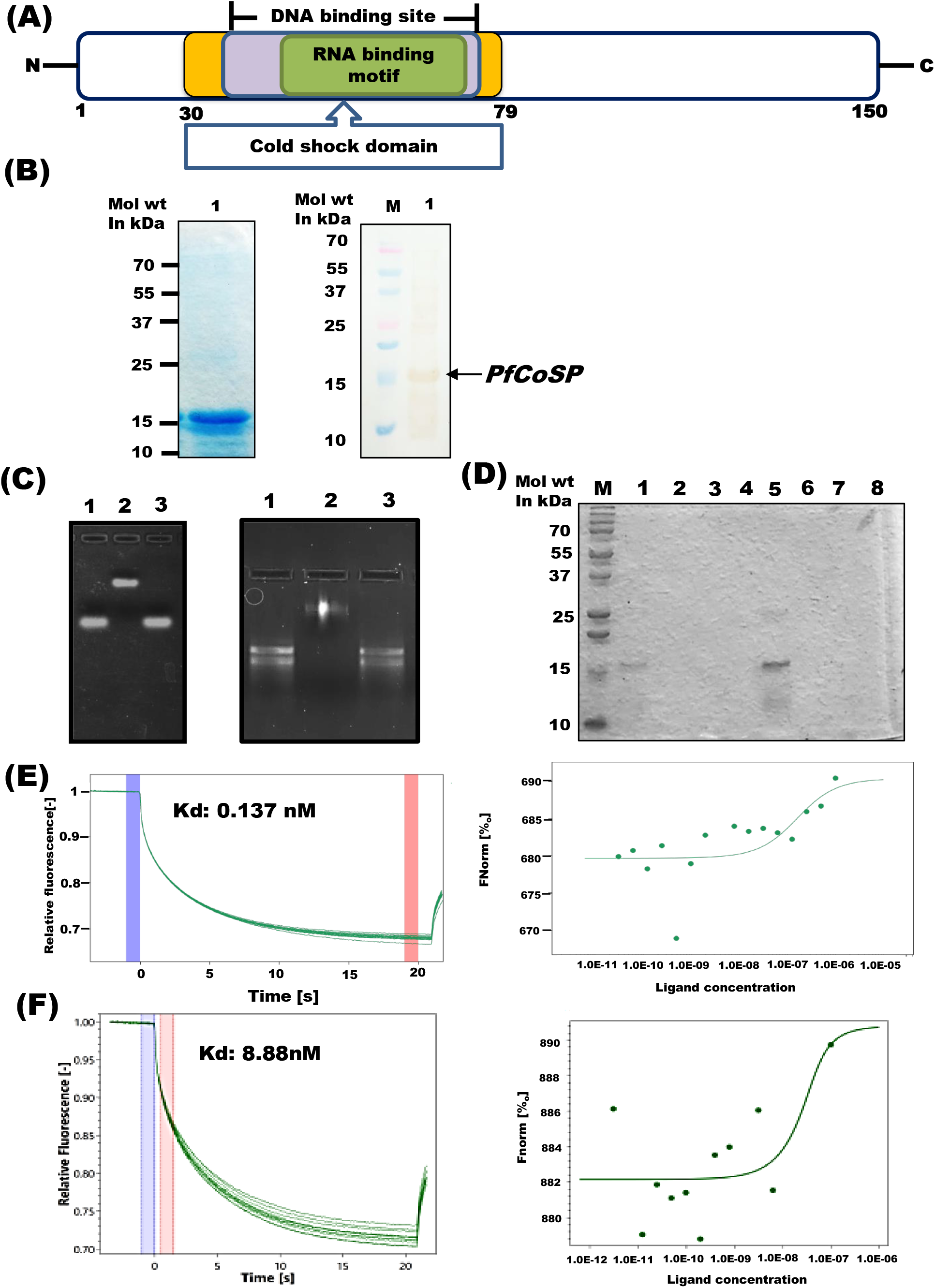
PfCoSP exhibits DNA and RNA binding ability. **(A)** Schematic representation of predicted domains and motifs of *Pf*CoSP. **(B)** (Left panel): SDS-PAGE showing recombinant purified *Pf*CoSP. Lane 1: *Pf*CoSP. (Right panel): Western blot analysis of recombinant purified *Pf*CoSP using anti-histidine antibodies. **(C)** Gel retardation assay showing *Pf*CoSP-nucleic acid interaction. Left panel: Agarose gel showing *Pf*CoSP-DNA binding. Lane 1: genomic DNA, lane 2: *Pf* genomic DNA + *Pf*CoSP, Lane 3: *Pf* genomic DNA + BSA. Right panel: Agarose gel showing *Pf*CoSP-*Pf*RNA binding. Lane 1: *Pf* RNA, lane 2: *Pf*RNA+ *Pf*CSP, Lane 3: *Pf*RNA + BSA. **(D)** *Pf*CoSP-DNA interaction using pull down assay. *Pf*CoSP and negative control BSA were incubated with single stranded (ss) DNA oligomer/double stranded (ds) DNA immobilized cellulose beads in a pull down assay. Elutes from assay were loaded on 15% SDS-PAGE. Lane L: Molecular weight marker, lane 1: ds DNA+ *Pf*CoSP, lane 2 : ds DNA wash, lane 3: ds DNA+ BSA, lane 4: BSA wash, lane 5, ss DNA + *Pf*CoSP, lane 6: ssDNA wash, lane 7: ssDNA+ BSA, lane 8: ss DNA wash. **(E, F)** *Interaction studies of Pf*CoSP *with Pfg*DNA (E) and *Pf*RNA (F) using Nano temper’s Monolith assay. Purified recombinant *Pf*CoSP was immobilized on gold sensor chip followed by titration with varying concentrations of *Pfg*DNA (E) and *Pf*RNA (F). Net dissociation graphs (left panels) generated K_d_ of 0.137nM and 8.88nM for *Pf*CoSP-gDNA (upper panel) and *Pf*CoSP-*Pf*RNA (lower panel) interactions respectively. Dose response curves for *Pf*CoSP-gDNA (upper panel) and *Pf*CoSP-*Pf*RNA (lower panel) are represented in right panel.

### *Pf*CoSP gives positive effect on growth of transformed *E. coli* cells upon cold stress

We investigated the effect of *Pf*CoSP on growth of BL21 (DE3) *E. coli* cells transformed with its cloned plasmid by subjecting them to low temperature. Transformed cells were incubated at 20°C post protein induction for 8 hours and samples were taken at 4 and 8 hrs of treatment. We observed that the growth of cells was more in case of transformed cells as compare to control where induction for *Pf*CoSP expression was not given (Fig. S1). This suggests that *Pf*CoSP has properties that give positive effect on growth of cells upon cold stress.

### *Pf*CoSP binds with nucleic acids

*Pf*CoSP carries DNA and RNA binding domains. Therefore, we used gel retardation assay and pull-down assays to test nucleic acid binding properties of *Pf*CoSP. *Pf*CoSP was incubated separately with *Pf* gDNA and RNA and subjected to agarose gel electrophoresis. Retardation in migration of nucleic acids was observed suggesting that *Pf*CoSP binds with both DNA (Fig. 1C, lane 2; left panel) and RNA (Fig. 1C, lane 2, right panel). Negative control where BSA was incubated with DNA and RNA did not show any shift in an agarose gel upon incubation with *Pf* gDNA (Fig. 1C, lane 3; left panel) and RNA (Fig. 1C, lane 3; right panel), highlighting the specificity of the assay. The nucleic acid binding property was also analysed using nucleic acid hybridization assay with cellulose beads containing immobilized calf thymus ssDNA and ds DNA. 1 μgm *Pf*CoSP was mixed with cellulose matrix and incubated at 4□ for 30 mints. Bead pellet was extensively washed and boiled in 1X SDS-PAGE sample loading dye before loading on 15% SDS-PAGE. Band for *Pf*CoSP was detected in boiled beads sample, indicative of its interaction with both ssDNA and ds DNA (Fig. 1D, lane 1, 5). No protein band was detective in negative control where beads were incubated with BSA (Fig. 1D, lane 3,7).

Kinetic analysis of one-to-one interaction of purified recombinant *Pf*CoSP with *Pf*gDNA and *Pf*RNA was carried out using NanoTemper Monolith NT.115 instrument. Fig. 1E and F shows association and dissociation curve for *Pf*CoSP-*Pf*gDNA and *Pf*CoSP-*Pf*RNA binding respectively. Steady-state analysis reveals the net disassociation constant, which is generated by plotting response at equilibrium as a function of unlabeled protein concentration (ranging from 250 nM to 5 µM). K_d_ for *Pf*CoSP*-Pf*g*DNA* and *Pf*CoSP-*Pf*RNA binding was observed to be 0.137 nM and 8.88 nM respectively.

### *Pf*CoSP interacts with cytoskeleton proteins alpha tubulin and beta tubulin

Al-Fageeh et. al. reported a proposed model for the co-ordinated cellular responses in mammalian cells upon exposure to mild hypothermia cold-shock. In the model, cold shock proteins can link transcription and translation via interactions with target mRNAs and the cytoskeleton [36]. In light of the above fact, we tested the binding of *Pf*CoSP with components of cell cytoskeleton *viz*. alpha and beta tubulin using *in vitro* and *in vivo* assays. Preliminary screening was performed using semi-quantitative ELISA assays that depict significant binding of *Pf*CoSP with alpha and beta tubulin in a concentration dependent manner. Binding of *Pf*CoSP with alpha and beta tubulin showed saturation at higher concentrations (Fig. 2A).

**Figure 2:**
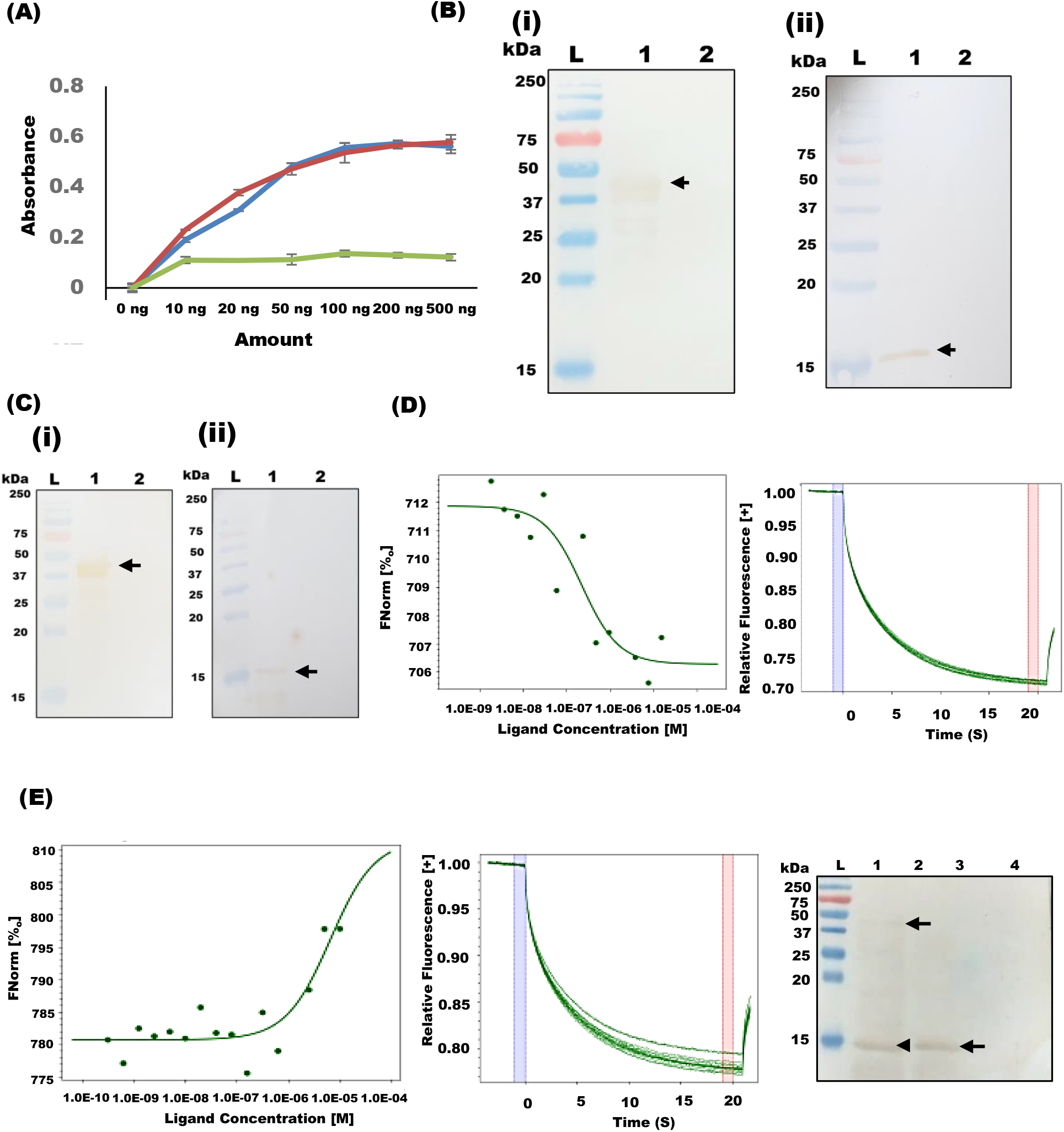
Interaction of PfCoSP with alpha and beta tubulin. **(A)** Semi-quantitative ELISA. Concentration-dependent binding curves of *Pf*CoSP with alpha and beta tubulin where y-axis represents absorbance at 490 nm and x-axis denotes amount of *Pf*CoSP. Error bars represent standard deviation among three replicates. Blue and red line depicts *Pf*CoSP-alpha tubulin and *Pf*CoSP-beta tubulin binding respectively whereas green depicts BSA as negative control. **(B, C)** Co-immunoprecipitation assay. *Pf*CoSP and alpha/beta tubulin specific antisera were coupled to aminolink plus coupling resin, and used to pull down alpha/beta tubulin and *Pf*CoSP respectively from parasite culture. (i) L: Ladder, 1: Elute from *Pf*CoSP specific antisera coupled beads probed with alpha tubulin antibodies, lane 2: Elute from preimmune sera coupled beads. (ii) L: Ladder, lane 1: Elute from alpha tubulin specific antisera coupled beads probed with *Pf*CoSP antibodies, lane 2: Elute from preimmune sera coupled beads. (Ci) Elute from *Pf*CoSP specific antisera coupled beads probed with beta tubulin antibodies, lane 2: Elute from preimmune sera coupled beads. ii) Elute from beta tubulin specific antisera coupled beads probed with *Pf*CoSP antibodies, lane 2: Elute from preimmune sera coupled beads. **(D, E)** *Interaction studies of PfCoSP with alpha tubulin (D) and beta tubulin (E) using Nano temper’s Monolith assay*. Purified recombinant alpha and beta tubulin were immobilized on gold sensor chip followed by titration with varying concentrations of *Pf*CoSP. Net dissociation graphs (left panels) generated K_d_ of 210 nM and 6.49 µM for *Pf*CoSP-alpha tubulin (D) and *Pf*CoSP-beta tubulin (E) interactions respectively. Dose response curves for *Pf*CoSP-alpha tubulin (D) and *Pf*CoSP-beta tubulin (E) are represented in right panel. **(F)** *Complex binding assays*. Complex formation of *Pf*CoSP with RNA and alpha tubulin. Lane L: Molecular weight marker, lane 1: Elute from *Pf*CoSP bound to single stranded (ss) DNA oligomer immobilized cellulose beads incubated with alpha tubulin, lane 2: Elute from single stranded (ss) DNA oligomer immobilized cellulose beads incubated with *Pf*CoSP (Positive control). lane 3: Elute from beads incubated with alpha tubulin (negative control). Lane 4: Elute from beads incubated with BSA followed by alpha tubulin (negative control).

*In vivo* co-immunoprecipitation assays followed by western blot analysis also validated the above results. *Pf*CoSP specific antisera was coupled to aminolink plus coupling resin, and used to pull down alpha and beta tubulin from parasite culture. Elutes from the assay were probed by western blot analysis using anti-alpha and beta tubulin antibodies separately. While a distinct band corresponding to alpha and beta tubulin eluted from *Pf*CoSP specific antisera coupled beads (Fig. 2 B, C lane 1; left panel), none was observed for preimmune sera linked beads (negative control) (Fig. 2B, C lane 2; left panel). Reverse co-immunoprecipitation assays further validated the interaction where anti-alpha and beta tubulin antibodies were coupled to aminolink plus coupling resin separately, and used to pull down *Pf*CoSP from parasite culture. Elutes from the assay were probed by western blot analysis using anti-*Pf*CoSP antibodies separately. A distinct band corresponding to *Pf*CoSP eluted from alpha and beta tubulin specific antisera coupled beads (Fig. 2B, C, lane 1; Right panel), while none was observed for preimmune sera linked beads (negative control) (Fig. 2B, C lane 2; right panel).

We also performed kinetic analysis of one-to-one interaction of purified recombinant *Pf*CoSP with alpha and beta tubulin using NanoTemper Monolith NT.115 instrument. We observed significant binding of *Pf*CoSP alpha and beta tubulin with. Fig. 2D C shows association and dissociation curve for *Pf*CoSP-alpha/beta tubulin binding. Steady-state analysis reveals the net disassociation constant, which is generated by plotting response at equilibrium as a function of unlabeled protein concentration (ranging from 0.0061 µM to 20 µM). K_d_ for *Pf*CoSP-alpha/beta tubulin binding was observed to be 210 nM and 6.49 µM respectively (Fig. 2D).

### *Pf*CoSP binds with RNA and tubulin simultaneously to form a complex

Since *Pf*CoSP binds to both RNA and alpha/beta tubulin, we tested its potential to attach with both simultaneously to form a complex using cellulose beads with immobilized single and double stranded calf thymus DNA. Recombinant *Pf*CoSP was incubated with the beads, and post washing allowed to bind with alpha/beta tubulin separately. Elutes from the assay were analysed by western blot analysis using specific polyclonal antisera against *Pf*CoSP and alpha/beta tubulin simultaneously. Distinct bands of *Pf*CoSP and alpha tubulin eluted from beads were observed (Fig. 2E, lane 1), while no band was detected where beads were first incubated with BSA followed by alpha tubulin (Fig. 2E, lane 2), showing the specificity of the assay. *Pf*CoSP was incubated to beads as positive control. These data suggest that *Pf*CoSP binds with RNA and alpha tubulin simultaneously to form a complex. No band for beta tubulin was observed from *Pf*CoSP bound RNA beads (Data not shown).

***Pf*CoSP is expressed in the asexual blood stages and gametocyte stage of parasite**

Expression of *Pf*CoSP was evaluated at transcript level during the asexual blood stages of *Pf*3D7. Our RT-PCT data showed that *Pf*CoSP expresses at all three stages of asexual blood stage of *Pf*3D7. However, the expression of *Pf*CoSP is many folds higher in trophozoite and schizont stages as compared to ring stage of malaria parasite (Fig. 3A). 18S rRNA was used as a loading control to clarify equal loading of cDNA sample.

**Figure 3:**
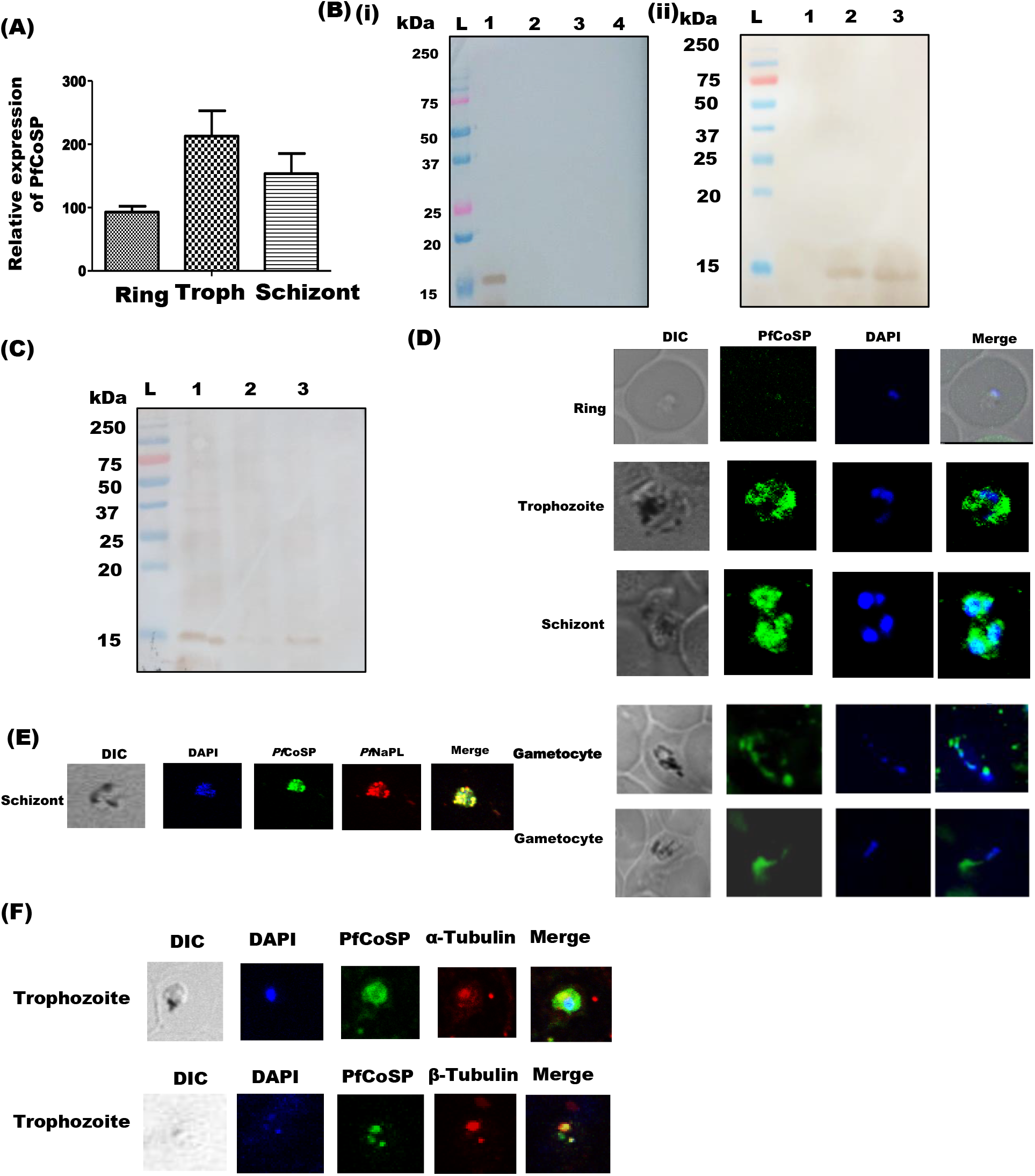
In vivo expression and localization of PfCoSP. **(A)** Relative expression of *Pf*CoSP at transcript level in asexual blood stages of malaria parasite using RT-PCR. Stages are labelled below the bars. **(B i)** Detection of *Pf*CoSP in infected erythrocytes by anti-*Pf*CoSP antibodies. Uninfected erythrocytes were loaded as negative control. Lane L: Molecular weight marker indicated in kDa, lane 1: Infected erythrocytes, lane 2: Infected erythrocytes cytosolic fraction, Lane 3: Uninfected erythrocytes, Lanes 4: Uninfected erythrocytes cytosolic fraction. **(B ii)** Western blot analysis of stage specific *Pf*3D7 parasite lysates to test *in vivo* expression of *Pf*CoSP. Lane L: protein ladder; lane 1: Ring stage parasite lysate. Lane 2: Trophozoite stage parasite lysate. Lane 3: Schizont stage parasite lysate. Blot was probed with anti-*Pf*CoSP antibodies followed by secondary antibodies. **(C)** Expression of *Pf*CoSP upregulates on cold stress. Schizont stage culture was treated at 25°C, 37 °C, 42 ° C for 6 hours and samples were subjected to western blot analysis followed by probing with anti-*Pf*CoSP antibodies. Lane L: Ladder, lane 1: parasite lysate treated at 25 °C, lane 2: parasite lysate treated at 42 °C, lane 3: parasite lysate treated at 37 °C. **(D)** Expression and localization analysis of *Pf*CoSP at different asexual and gametocyte stages of parasite life cycle. Smears of methanol-fixed *Pf*3D7-infected erythrocytes were stained with anti-*Pf*CoSP antibodies (1:250) followed by incubation with Alexa Fluor-conjugated secondary antibodies (Alexa Fluor 546, green). DIC: differential interference contrast image, DAPI: nuclear staining 40, 6-diamidino-2-phenylindole (blue); *Pf*CoSP: mouse anti-*Pf*CoSP (green); merge: overlay of *Pf*CoSP with DAPI. **(E)** Co-localization of *Pf*CoSP with *Pf*NapL. Smears of methanol-fixed *Pf*3D7-infected erythrocytes were stained with anti-*Pf*CoSP (1:250) and anti-*Pf*NapL antibodies (1:250), followed by incubation with Alexa Fluor-conjugated secondary antibodies (Alexa Fluor 488, red; Alexa Fluor 546, green). DIC: differential interference contrast image, DAPI: nuclear staining 40, 6-diamidino-2-phenylindole (blue); *Pf*CoSP: mouse anti-*Pf*CoSP (green); *Pf*NapL-: anti-*Pf*NapL antibody (red); merge: overlay of *Pf*CoSP with *Pf*NapL. **(F)** Co-localization of *Pf*CoSP with tubulin. Smears of methanol-fixed *Pf*3D7-infected erythrocytes were stained with anti-*Pf*CoSP (1:250) and anti-tubulin antibodies (1:250), followed by incubation with Alexa Fluor-conjugated secondary antibodies (Alexa Fluor 488, red; Alexa Fluor 546, green). DIC: differential interference contrast image, DAPI: nuclear staining 40, 6-diamidino-2-phenylindole (blue); *Pf*CoSP: mouse anti-*Pf*CoSP (green); *Pf*Tubulin-: anti-*Pf*tubulin antibody (red); merge: overlay of *Pf*CoSP with *Pf*tubulin.

Expression analysis of *Pf*CoSP during the asexual blood stages was investigated using specific antisera against *Pf*CoSP. Western blot analysis on total protein extracts of mixed stage *Pf* 3D7 infected RBCs detected a single protein band at ∼ 17 kDa (Fig. 3B, lane 1; left panel,) for *Pf*CoSP. An identical blot probed with pre-immune sera showed no signal (Fig. S2 left panel). Uninfected erythrocytes lysates were used as negative control in the experiment. The expression of the *Pf*CoSP protein was also explored with different asexual stages of the parasite by western blotting using anti-mice *Pf*CoSP antibodies. *Pf*CoSP was detected in trophozoites (27 to 33 hpi) and schizonts (42 to 48 hpi) but not in ring-stage parasites (15 to 21 hpi) (Fig. 3C; left panel). A parallel western blot assay was also developed with tubulin specific antibodies as a loading control for the parasite proteins (Fig. S2, middle panel).

We also investigated the change in expression of *Pf*CoSP on giving temperature treatment in malaria parasite by subjecting schizont stage cultures to 25°C for 6 hours. Samples were subjected to western blot and probed with anti-*Pf*CoSP antibodies. Expression of *Pf*CoSP was observed many folds higher at 25°C as compared to 37°C and 42°C (Fig. 2C), suggesting its significant role in malaria parasite on cold stress. A parallel western blot was also developed with tubulin specific antibodies as a loading control for the parasite proteins (Fig. S2, right panel).

The expression of *Pf*CoSP during the asexual blood stages of *Pf* 3D7 and gametocyte stages of *Pf* RKL9 were investigated by indirect immunofluorescence assays using anti-*Pf*CoSP antibodies. Thin blood smears of mixed stage *Pf* 3D7 and *Pf* RKL9 cultures were fixed with methanol and blocked with 5% BSA in PBS. The slides were probed with anti-*Pf*CoSP antibodies (1:200) followed by Alexa Fluor 546 conjugated anti-mice secondary antibodies (1:200). The parasite nuclei were counterstained with DAPI (4′,6′-diamidino-2-phenylindole). Our data showed that *Pf*CoSP is expressed at asexual blood stages as well as gametocyte stages of the parasite (Fig. 3E). Expression of *Pf*CoSP was found to be limited to trophozoites and schizonts but not in ring-stage parasites (Fig. 3D). This data was consistent with our stage specific detection of *Pf*CoSP in parasite by western blotting. Fluorescence pattern observed for *Pf*CoSP at asexual blood stages was suggestive of its localization to parasite nucleus and cytosol. To confirm its cytosolic localization, co-localization of *Pf*CoSP was performed with a reported cytosolic protein, *Pf*NapL. *Pf*CoSP showed significant co-localization with *Pf*NapL that further confirmed its parasite cytosolic localization (Fig. 3E).

We also performed co-localization of *Pf*CoSP with alpha/beta tubulin during the asexual stages of *Pf* using protein-specific antibodies on cultured parasites. Our data showed significant overlapping of signals for *Pf*CoSP and alpha/beta tubulin, suggestive of their colocalization (Fig. F). These results further support interaction of *Pf*CoSP with alpha and beta tubulin.

### ‘LI71’ binds with *Pf*CoSP and inhibits *Pf*CSP -alpha/beta tubulin interactions

We performed extensive literature survey to look for an inhibitor for a cold shock protein. We found a molecule named L171 that directly binds the cold shock domain of a human cold shock protein ‘LIN28’ to suppress its activity against let-7 in leukemia cells and embryonic stem cells [37]. Multiple sequence alignment of *Pf*CoSP with LIN28 suggests significant similarity between the proteins (Fig. 4A). Therefore, we tested this inhibitor for binding with *Pf*CoSP and inhibiting its interaction with alpha/beta tubulin. NMR spectra of LI71 is represented in Fig. S3, S4, S5, S6 and its synthetic scheme is represented in Fig. S7. Preliminary analysis of interaction was performed by docking modeled structure of cold shock domain of *Pf*CoSP with LI71 using Patchdock [38]. Modeling details of *Pf*CoSP are described in our previous report [39]. We observed that LI71 binds well into the cavity of *Pf*CoSP with a binding energy of -64.01 KJ/mol (Fig. 4B, left panel). Analysis of docked structure using Prankweb webserver [40] gave detail analysis of binding site. Residues of *Pf*CoSP forming the pocket for LI71 and involved in interaction include I46, V57, H58, Y59, T60, D61, R66, T67, F68, A79, W80 and N81. Out of these, we have shown earlier that H58 and Y59 forms maximum number of interactions with BDNA and RNA [39].

**Figure 4:**
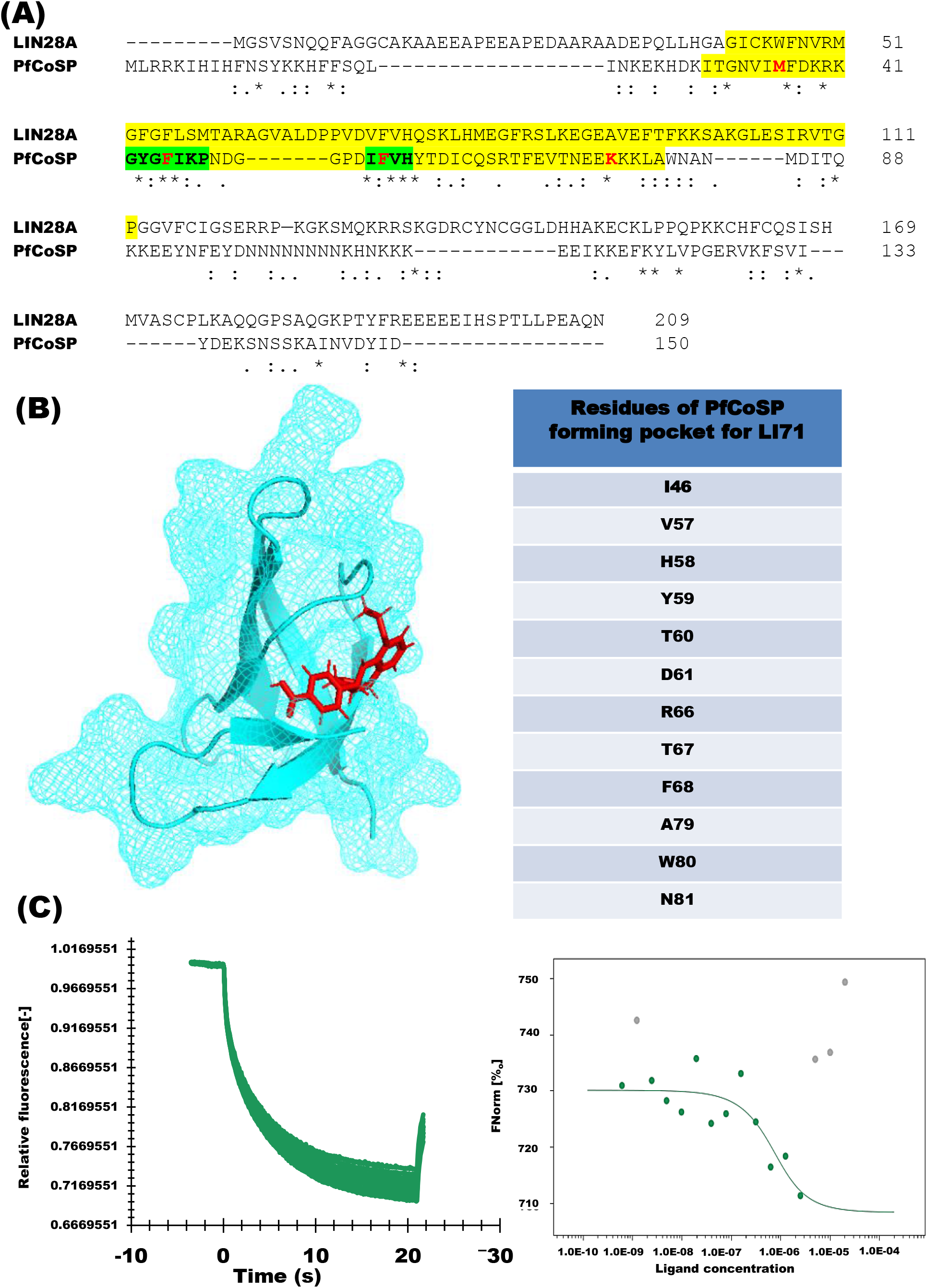
Binding of PfCoSP with LI71. **(A)** Multiple sequence alignment of *Pf*CoSP with LIN28A. Residues highlighted in yellow constitute cold shock domain. Residues marked with red are predicted residues involved in DNA binding and those highlighted in green are involved RNA binding. **(B)** Docking of *PfCoSP with LI71*. Left panel: Structural representation of docked *Pf*CoSP-LI71 complex. Cyan represents *Pf*CoSP whereas red denotes LI71. Right panel: Residues of *Pf*CoSP predicted to form the pocket for LI71 binding are tabulated. **(C)** Interaction studies of *Pf*CoSP with LI71 using Nano temper’s Monolith assay. Purified recombinant *Pf*CoSP was immobilized followed by titration with varying concentrations of LI71. Net dissociation graph (left panel) generated K_d_ of 431nM for *Pf*CoSP-LI71 interaction. Dose response curves are represented in right panel.

**Figure 5:**
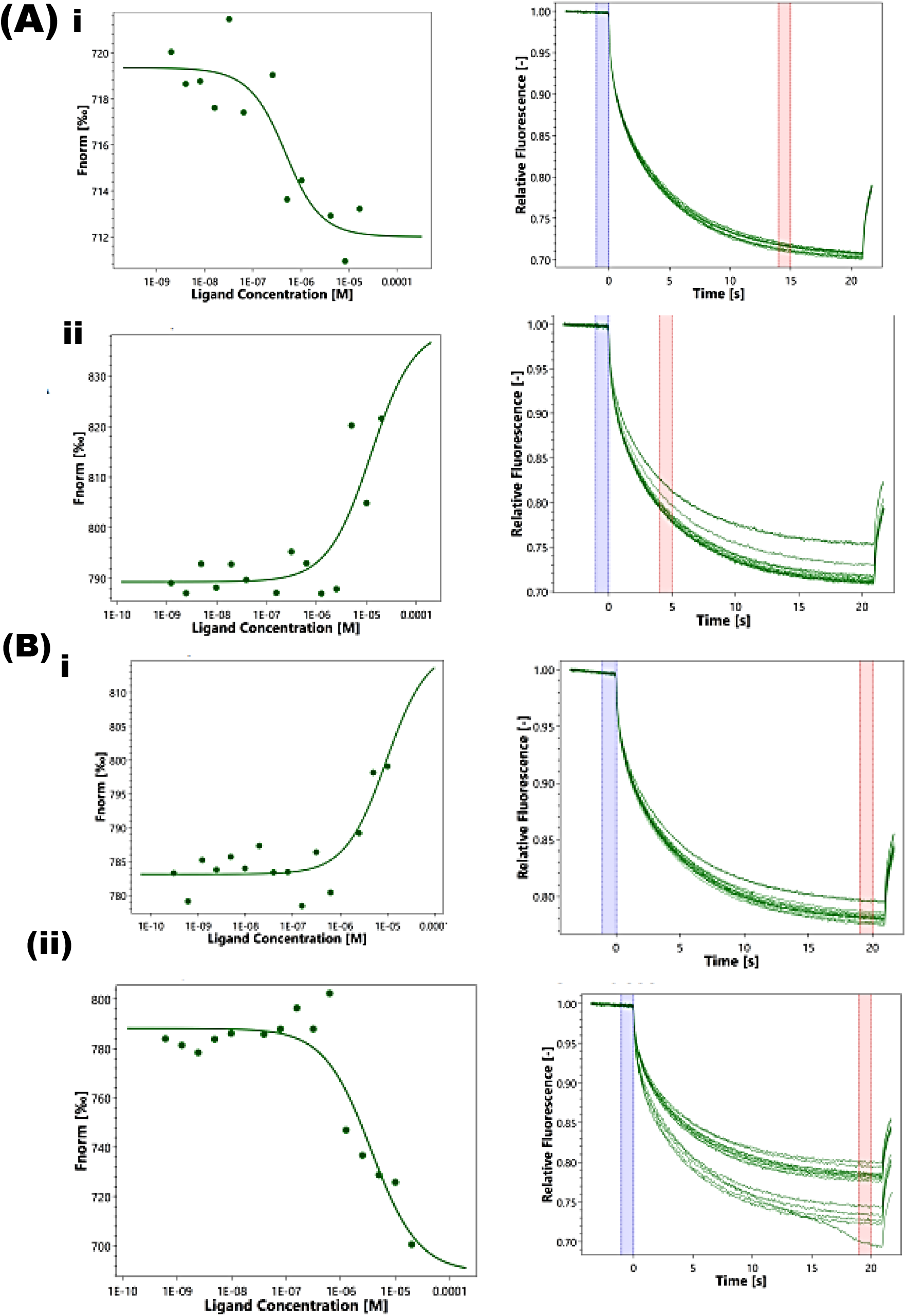
LI71 competes with α-tubulin and β-tubulin to interact with *Pf*CoSP. **(A)** Interaction of α-tubulin with recombinant *Pf*CoSP NHS-Red dye labelled protein (i) was evaluated through MST. The K*d* constant of 274 ± 64 nM was obtained in case of interaction of α-tubulin with *Pf*CoSP. A decreasing MST signal (F_norm_ [%□] starting at 712 units, decreasing to 706 units) was observed with increasing concentration of *Pf*CoSP yielding a sigmoidal curve. (ii) Competition with compound L171 as a competing molecule interfering with the interaction of α-tubulin-CoSP displayed F_norm_ level at 790 units increasing to 835 units with K*d* 12.65 ± 0.45 µM with increasing concentration of the compound. **(B)** Similarly, interaction between β-tubulin with CoSP (i) was also analysed. Labelled β-tubulin was titrated against serially diluted *Pf*CoSP. β-tubulin also displayed interaction with *Pf*CoSP with K*d* value 7.92 ± 1.43 µM with F_norm_ starting from 785 units and increasing to 810 units yielding a sigmoidal dose-response curve. When the interaction between β-tubulin and *Pf*CoSP was interrupted by increasing concentrations of compound L171, F_norm_ value decreased from 785 to 710 units, yielding a K*d* value of 3.055 ± 0.545 µM (ii).

Kinetic analysis of one-to-one interaction of purified recombinant *Pf*CoSP with L171 was carried out using NanoTemper Monolith NT.115 instrument. We observed significant binding of *Pf*CoSP with L171. Fig. 4 C shows association and dissociation curve for *Pf*CoSP-L171 binding. Steady-state analysis reveals the net disassociation constant, which is generated by plotting response at equilibrium as a function of unlabeled protein concentration (ranging from 0.0061 µM to 20 µM). K_d_ for *Pf*CoSP*-*LI71 binding was observed to be 431 nM.

MST can be employed to perform competition experiment owing to several advantages it poses. The technique is label-free and site-specific resulting in reliable information regarding the ability of a compound to interrupt the protein-protein interaction (Jerabek-Willemsen et al., 2011). Labelled α-tubulin was titrated against serially diluted *Pf*CoSP followed by analyses with Monolith NT.115 instrument. In case of interaction of α-tubulin with *Pf*CoSP, a decreasing MST signal (F_norm_ [%[] starting at 712 units, decreasing to 706 units) was observed with increasing concentration of *Pf*CoSP yielding a sigmoidal curve giving a K*d* value of 274 ± 64 nM. This low K*d* value represents a strong interaction between α-tubulin with *Pf*CoSP. However, the competition experiment performed with compound LI71 as a competing molecule interfering the interaction of α-tubulin-*Pf*CoSP displayed F_norm_ level at 790 units increasing to 835 units with K_d_ 12.65 ± 0.45 µM while increasing concentration of the compound. This suggests a significant binding of LI71 with *Pf*CoSP displacing the labelled α-tubulin based on the MST signals.

Similarly, interaction of β-tubulin with *Pf*CoSP in the presence of LI71 was also analysed. Concentration of labelled β-tubulin was kept constant and titrated against serially diluted *Pf*CoSP. On observation, β-tubulin also displayed interaction with *Pf*CoSP with K_d_ value 7.92 ± 1.43 µM with F_norm_ starting from 785 units and increasing to 810 units. When the interaction between β-tubulin and *Pf*CoSP was interrupted by increasing concentrations of compound LI71, F_norm_ value decreased from 785 to 710 units, yielding a K_d_ value of 3.055 ± 0.545 µM. Thus, this change in the MST signal suggests competing property of LI71 against β-tubulin to interact with *Pf*CoSP. Overall, based on our findings we can interpret L171 as a compound competing with both the tubulins to bind with CoSP.

### LI71 inhibits *in vitro* growth of malaria parasites

LI71 was screened for its antimalarial activity against asexual stage of human malaria parasite *in vitro*. For an IC_50_ determination, highly synchronized ring-stage parasites of 3D7 and RKL-9 were exposed to a range of compound concentration for 72 h. Parasitaemia was measured at 72 h post-treatment using SYBR Green I based fluorescence assay and graph was plotted for the average value of three independent sets of experiments. Our data revealed potent anti-malarial activity of LI71 against 3D7 and RKL strain of *P. falciparum* with a significant reduction in the parasite load. IC_50_ value of LI71 in 3D7 and RKL strain of *P. falciparum* was found to be 1.35 nM and 6.3 nM respectively (Fig. 6A).

**Figure 6:**
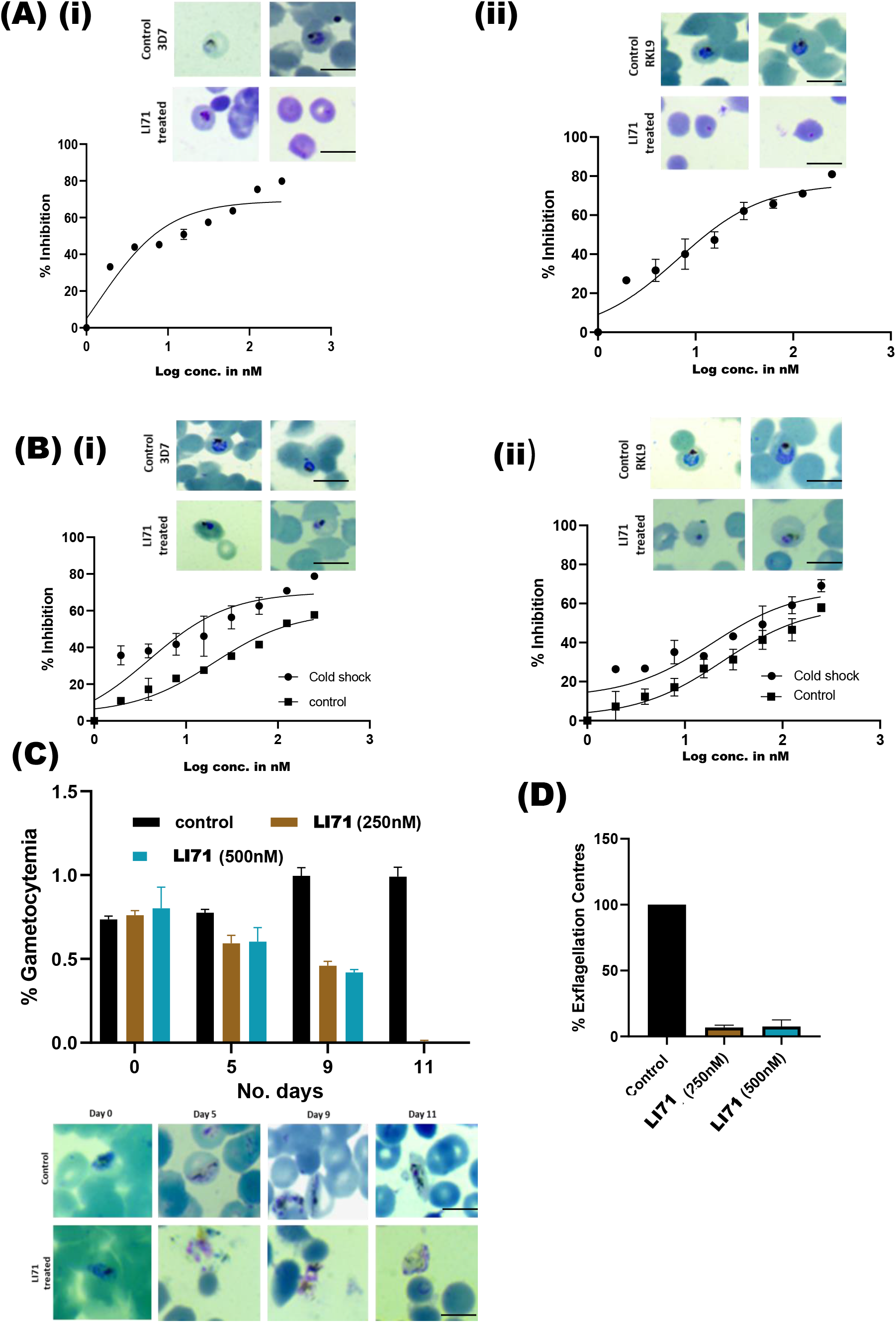
Growth inhibitory effect of LI71 on Pf3D7. **(A)** The parasite culture synchronized at ring stage was treated with different concentrations (250, 125, 62.5, 31.25, 15.62, 7.8, 3.9, 1.95 nM) of LI71 for 72 h, and the percent growth inhibition was estimated. IC_50_ values of LI71 in *Pf*3D7 (i) and *Pf*RKL-9 (ii) were evaluated by plotting growth inhibition values against the log concentration of LI71. The experiment was done in triplicate, and the results were shown as mean values ± SD. **(B)** Anti-malarial activity of LI71 on *Pf*3D7 (i) and *Pf*RKL-9 strain (ii) at low temperature. Parasite was treated with LI71 for 4 hours at 19°C. Post washing, the culture was kept at 37°C for 68 hrs. As a control, parasite was treated with LI71 for 4 hours at 37°C followed by a wash. IC_50_ values of LI71 were evaluated by plotting growth inhibition values against the log concentration of LI71. The experiment was done in triplicate, and the results were shown as mean values ± SD. **(C)** LI71 inhibit maturation of *Pf* gametocytes. (i) Bar graph represents number of gametocyte determined by counting Giemsa-stained smears (relative to DMSO control on day 1,5,9,11) after treatment with LI71 at concentration 250 nM and 500 nM. Error bars represent mean[±[S.D with P-value < 0.05 * **(D)** LI71 inhibit ex-flagellation of male gametocyte. Late stage *P. falciparum* gametocyte was mixed with pre-warmed complete RPMI. and incubated with LI71 at concentration 250 nM and 500 nM for 30 min at 37°C. Ex-flagellation was induced by transferring the sample to 25°C in the presence of 100 µM of xanthurenic for 15 min. Sample were then dispensed onto glass bottom plate and observed under bright field microscopy at 40X magnification. Bar graphs represent the no of ex-flagellation centres observed in 10 field views post addition of LI71 compound. Data represent the mean ±[S.D (n=3)

### Thermotolerance in malaria parasite requires *Pf*CoSP

Our data showed that LI71 binds with *Pf*CoSP and inhibits its interaction with its binding partner tubulin. Also, LI71 showed significant anti-malarial activity. In light of above facts, we presume that LI71 inhibits the function of *Pf*CoSP *in vivo*. Since cold shock proteins play an important role in living organisms to survive under cold conditions, we hypothesize that expression of *Pf*CoSP is also required for parasite growth during temperature stress. We tested this hypothesis by inhibiting *Pf*CoSP by LI71 to know whether its inhibition influence parasite growth upon cold (21°C) stress. Both 3D7 and RKL-9 parasites were treated with inhibitor for 4 h at 21°C followed by 37°C for 72 h. SYBR Green I analysis was performed to determine the IC50 in stressed condition. Our data showed that inhibition of *Pf*CoSP by treating parasites with LI71 significantly reduced growth after temperature stress in comparison to control (parasite treated with LI71 at 37°C) (Fig. 6B). Also, the IC50 under temperature stress condition was found to be lower in comparison to control (Fig. 6B). These data suggest that temperature stress render the parasite more susceptible to *Pf*CoSP inhibitor.

### LI71 effects the process of gametocytogenesis

LI71 was also tested for its effect on gametocytogenesis in *P. falciparum* gametocyte. RKL-9 strain of *P. falciparum* was differentiated into sexual stages and treated with L171 at 250 and 500 nM concentration. To identify when LI71 cause arrest, we performed microscopy of blood smears and quantified gametocyte stages. Our results suggest that daily dose of LI71 at early stage II cause the decline in gametocytemia (Fig. 6C). Giemsa image analysis revealed that in treated culture parasite has altered morphology while DMSO treated control gametocyte develop till stage V (Fig. 6C).

### LI71 inhibits the male gamete ex-flagellation *in vitro*

We carried out *in vitro* exflagellation assay with *P. falciparum* gametocyte to assess the effect LI71 on male ex-flagellation. Stage V *P. falciparum* gametocyte was pre-incubated with LI71 for 1 h at concentration of 250 & 500 nM at 37°C. Post incubation, ex-flagellation was induced by the addition of 100 µM xanthurenic acid at RT for 15 min. Afterwards, the number of ex-flagellation centers were microscopically counted in both DMSO and compound treated parasites at 40X magnification. Ex-flagellation is a highly time-dependent and characterized by the emergence of the male gametes from the infected erythrocyte after a period of approximately 15 to 20 minutes. Once they emerge from the gametocyte residual body, they migrate away from the ex-flagellation centre through flagellar locomotion and continue to move until either they reach a female gamete and fertilize it. It is easy to distinguish male gametes by light microscopy since they are highly motile cells. In a monolayer of erythrocytes, ex-flagellation causes local disturbance of nearby cells while leaving distant cells motionless. Treatment with LI71 showed a significant reduction in the numbers of exflagellation centers by approximately 90% in comparison to DMSO control (Fig. 6D). These data suggest that LI71 inhibits the male gamete ex-flagellation and implicate the significant role of *Pf*CoSP during exflagellation.

## Discussion

Cold shock proteins are believed to counteract the harmful effects of low temperature by inhibiting the formation of secondary structures in mRNA, thereby facilitating the process of translation [3, 6]. Members of cold shock proteins are well characterized in bacteria, plants and humans and are known for their ability to perform pleiotropic functions inside the cell [39]. In the present study, we have attempted to delineate the function of a novel cold shock protein (*Pf*CoSP) of malaria parasite. Understanding the molecular interplay of *Pf* cold shock protein provide functional insights into malaria biology.

Full length construct of *Pf*CoSP was cloned in pET-28a(+) vector and overexpressed in soluble form in a bacterial expression system [BL21 (DE3) *E. coli* cells]. Polyclonal antibodies were generated against *Pf*CoSP and found to be specific for the protein. Since cold shock proteins are known to provide cold tolerance, we first tested the ability of *Pf*CoSP to give any growth advantage to *E. coli* cells upon cold treatment. Our results demonstrate that overexpression of *Pf*CoSP enhanced the BL21 (DE3) *E. coli* growth at hypothermic cold conditions. These data gave us the hint for the functional role of *Pf*CoSP in malaria parasite when it enters the mosquito host and face low temperature. Cold shock proteins are characterized by the presence of cold shock domain that exhibits nucleic acid binding properties [3, 6], therefore we tested *Pf*CoSP for DNA/RNA binding ability using *in vitro* assays. Our results clearly demonstrate that *Pf*CoSP binds with both DNA and RNA, suggesting that protein is likely to function as nuclei acid chaperone *in vivo*. Several studies have reported both DNA and RNA binding activity of cold shock proteins including YB-1 of humans, WCSP1 of wheat, AtCSP1-CSP4 of *Arabidopsis*, OsCSP1 (Os02g0121100) and OsCSP2 (Os08g0129200) of rice [39]. By nucleic acid binding ability, cold shock proteins are involved in several DNA and mRNA dependent processes including DNA replication and repair, mRNA splicing and translation, thereby regulating gene expression [38]. Considering this, we suggest that *Pf*CoSP may play a significant role in regulating gene expression in malaria parasite.

It has been suggested that cytoskeleton disassembly upon cold shock disrupt translational machinery which in turn inhibit protein translation at low temperatures [36]. B. al-Fageeh et. al. proposed that cold shock proteins link transcription and translation via interactions with target mRNAs and the cytoskeleton [36]. In light of above facts, we tested the binding of *Pf*CoSP with components of cell cytoskeleton *viz*. alpha and beta tubulin. Our *in vitro* binding assays (semi-quantitative ELISA, co-immunoprecipitation assays and SPR) clearly illustrate for the first time that recombinant *Pf*CoSP binds to both alpha and beta tubulin in a concentration dependent manner. These data suggest the ability of *Pf*CoSP as a cytoskeleton linking protein.

Since *Pf*CoSP binds to both RNA and alpha/beta tubulin, we tested whether RNA and alpha/beta tubulin binds with *Pf*CoSP together to form a complex, or compete with each other for attachment to *Pf*CoSP. Our assays clearly show that *Pf*CoSP binds to both RNA and alpha tubulin simultaneously *in vitro* suggestive of separate binding sites for these on *Pf*CoSP. This highlights the ability of *Pf*CoSP to dually function as a nuclei acid chaperone and cytoskeleton linking molecule simultaneously.

The study was further sought to analyse the *in vivo* expression and localization of *Pf*CoSP at asexual blood stages and gametocyte stage of malaria parasite. *In vivo* expression of the protein is clearly evident from our western blot analysis on mixed stage parasite lysates which shows a single band at the expected size. Also, we observed that *Pf*CoSP expression upregulates many folds higher when parasite was subjected to cold conditions, further highlighting its role during cold stress. Immunofluorescence data depict the expression of *Pf*CoSP during gametocyte and asexual blood stages of malaria parasite. Stage specific expression analysis suggest expression of *Pf*CoSP mainly in the trophozoite and schizont stages. Also, our immunofluorescence data depict *Pf*CoSP as a nucleo-cytoplasmic protein. Further, *Pf*CoSP showed significant co-localization with *Pf* alpha and beta tubulin that suggest the interaction between the proteins.

We next performed extensive literature search to identify any inhibitor for a cold shock protein. A molecule named L171 was identified that directly binds the cold shock domain of a human cold shock protein ‘LIN28’ [37]. By interacting with CSD, LI71 suppresses activity of LIN28 against let-7 in leukemia cells and embryonic stem cells [37]. Multiple sequence alignment of *Pf*CoSP with LIN28 suggest significant similarity between the proteins. Our docking studies suggest that LI71 binds well into the cavity formed by cold shock domain of *Pf*CoSP. Therefore, we went further to test this inhibitor for binding with *Pf*CoSP and inhibiting its interaction with alpha/beta tubulin. Kinetic analysis of one-to-one interaction using NanoTemper Monolith NT.115 instrument suggested significant binding of *Pf*CoSP with L171 with a K_d_ of 431 nM. Our competitive inhibition assays using NanoTemper Monolith NT.115 instrument also suggested that LI71 inhibited *Pf*CoSP - alpha/beta tubulin interactions. These data provided a clue for the potential of this compound to obstruct the functions of *Pf*CoSP and prompted us to test the anti-malarial activity of LI71 on different stages of malaria parasite. We performed growth inhibition assays which clearly demonstrate that LI71 inhibited growth of blood stage *P. falciparum* 3D7 and chloroquine drug-resistant strain *Pf*RKL-9 with an IC_50_ value of 1.35 nM and 6.32 nM respectively. IC_50_ of LI71 against *Pf*3D7 and *Pf*RKL-9 was found to be even lower at cold stress. Since *Pf*CoSP is expressed at gametocyte stages, we studied cross-stage inhibitory potential of LI71. Our data revealed that LI71 attenuate gametocyte development and inhibit the male gamete ex-flagellation *in vitro*. This complementary activity of LI71 on the sexual stages of the parasites suggests its potential to reduce malaria transmission apart from its plausible role in chemoprevention. Overall, these results demonstrate antimalarial activity of LI71 against gametocytes and asexual blood stages of parasite.

Overall, our data highlights the potential of *Pf*CoSP that it may help the parasite growth and development upon cold stress when it enters mosquito host to complete its sexual cycle. In infected humans, majority of circulating parasites are asexually dividing merozoites. A small portion of these undergo a differentiation pathway to form sexually competent parasite called ‘gametocyte’. These highly specialized gametocytes are transmitted from an infected human to a susceptible mosquito host. *As* gametocytes switch from human host to female *Anopheles*, they egress from the host erythrocytes and face low temperature environment. We propose that during this transitioning, *Pf*CoSP play its pivotal role in adapting the parasite to cold stress condition.

It has been suggested that cytoskeleton disassembly upon cold shock disrupt translational machinery and might account for the suppression of protein synthesis in mammalian cells [36]. Based on our study, we propose a model for the role of *Pf*CoSP in assisting the process of translation by simultaneously interacting with mRNA targets and components of cell cytoskeleton (Fig. 7). In the model, *Pf*CoSP can capture mRNA targets as they emerge from the nucleus, or in the cytoplasm, and enable their maintenance of single stranded state to pursue efficient transcription and translation. *Pf*CoSP via interactions with the components of cell cytoskeleton also helps in forming a work bench for translation. In this way, *Pf*CoSP may play a significant role in regulating gene expression at different stages of malaria parasite. Since *Pf*CoSP seems essential for parasite survival, characterization of its interaction with DNA/RNA, tubulin and LI71 may form the basis for development of future anti-malarials.

**Figure 7:**
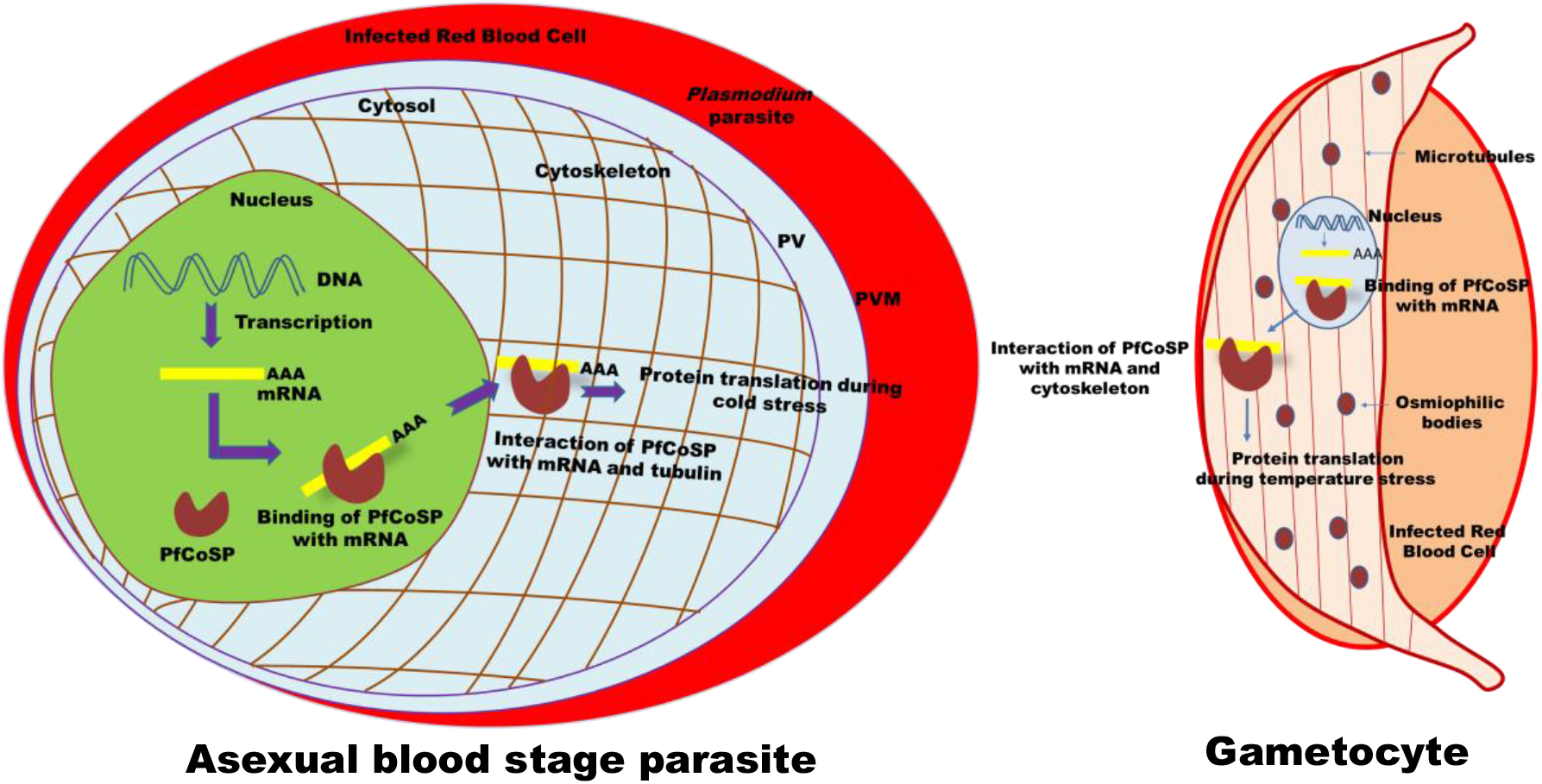
Probable functional role of *Pf*CoSP at gametocyte and asexual blood stages of malaria parasite. Upon temperature stress, *Pf*CoSP may bind mRNA targets and enable their maintenance of single stranded state to pursue efficient transcription and translation. In this way, *Pf*CoSP may play a significant role in regulating gene expression at different stages of malaria parasite.

## Acknowledgements

**AB** is supported by National Post-doctoral fellowship, SERB, India (Fellowship reference no. PDF/2019/000334). **MS** is supported by SNU-Foundation fellowship. **GK** is CSIR-SRF. **VK** was supported by Research Associateship Program of Department of Biotechnology, Govt. of India and **RS** is UGC-SRF, Govt. of India. Funding from Intensification of Research in High Priority Areas (IRHPA) of Science and Engineering Research Board (SERB) and the National Bioscience Award from DBT for SS is acknowledged.

## Declaration of interests

Authors declare no conflict of interests.

## Author contributions

AB: Experimental design, experimentation, data analysis and manuscript writing. MS: conducted all binding assays of LI71 using nanotemper and manuscript preparation. FDL: synthesized LI71. GK: conducted all experiments related to antimalarial activity of LI71 and manuscript preparation. VK: Experimental design, data analysis, conducted some binding assays using nanotemper and manuscript preparation. RS: conducted some binding assays using nanotemper. HS: Preparation of buffers, SG: Data analysis, CA: Syntheses of LI71, Data analysis and manuscript writing. SS: Conception of idea, experimental design, data analysis and manuscript writing.

## Notes

### Competing Interest Statement

The authors have declared no competing interest.

