## supplementary material for "Human malaria parasite cold shock protein plays an essential role in asexual and sexual stage development and presents an excellent druggable target"

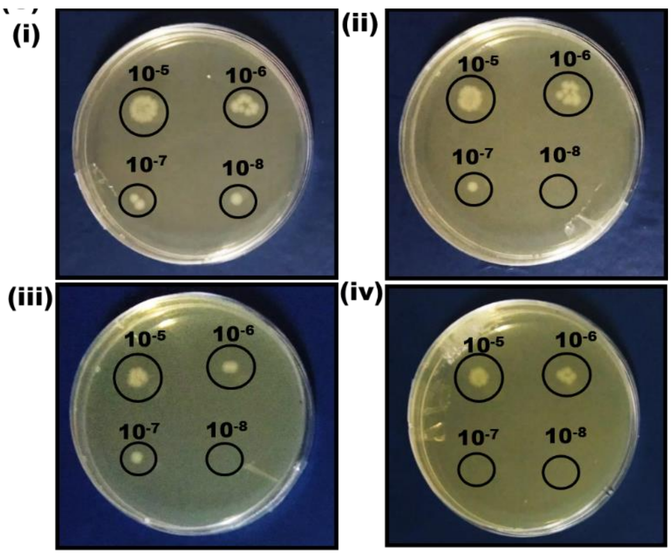


**Fig. S1:** Growth of *E.coli* cells transformed with cloned plasmid of *Pf*CoSP upon cold stress. (i, ii left panel) Colonies obtained after 4 hrs and 8 hrs of cold treatment. (i, ii right panel) Control plates after 4 and 8 hrs where induction was not given for *Pf*CoSP expression. Different dilutions used in the experiment are mentioned above the encircled colonies.

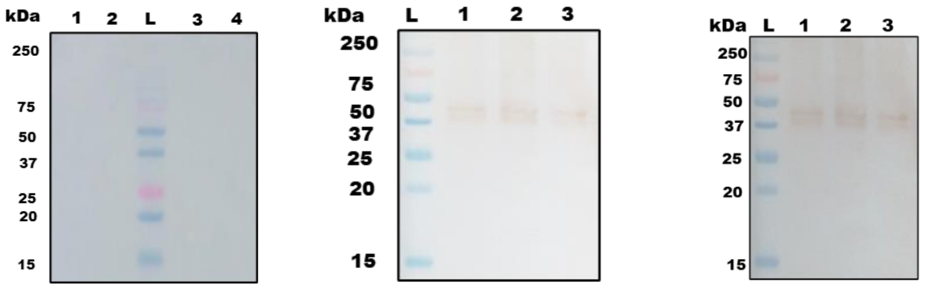


**Fig. S2:** Left panel represents control blot probed with pre-immune sera. Lanes L: Molecular weight marker indicated in kDa, lane 1: Infected erythrocytes, lane 2: Infected erythrocytes cytosolic fraction, Lane 3: Uninfected erythrocytes, Lanes 4: Uninfected erythrocytes cytosolic fraction. Middle blot represents western blot analysis with tubulin specific antibodies as a loading control for the parasite proteins. Lane L: protein ladder; lane 1: Ring stage parasite lysate. Lane 2: Trophozoite stage parasite lysate. Lane 3: Schizont stage parasite lysate. Right panel represent western blot developed with tubulin specific antibodies as a loading control for the parasite proteins. Schizont stage culture was treated at 25°C, 37 °C, 42 ° C for 6 hours and samples were subjected to western blot analysis followed by probing with anti-tubulin antibodies. Lane L: Ladder, lane 1: parasite lysate treated at 25 °C, lane 2: parasite lysate treated at 42 °C, lane 3: parasite lysate treated at 37 °C.

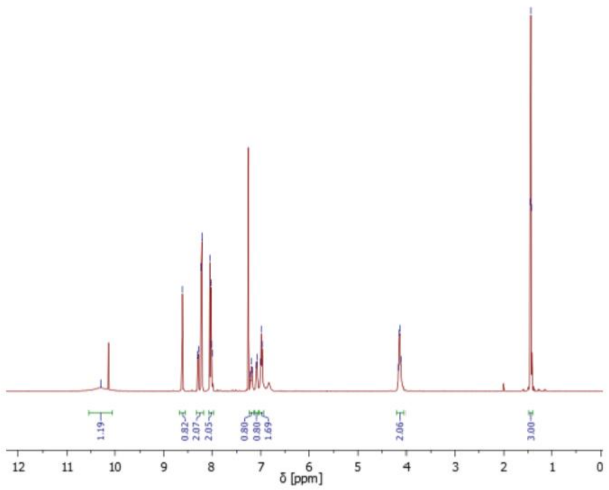


Fig. S3: $^{1}$H NMR spectrum of LI71 Precursor (Imine) (2)

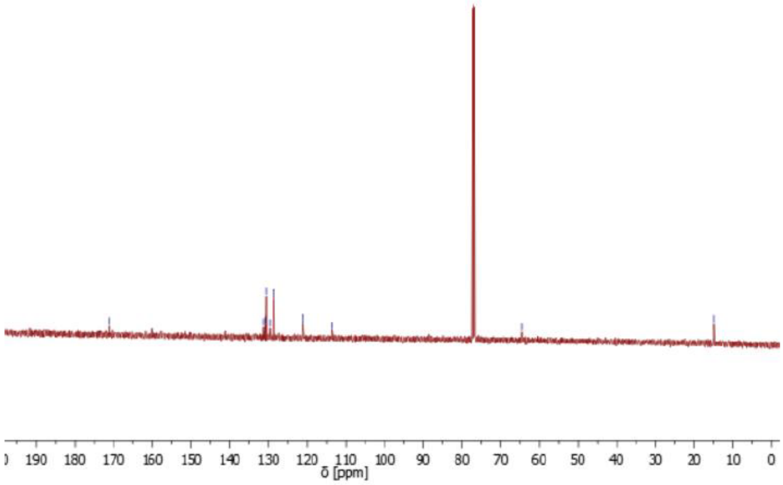


Fig. S4: $^{13}$C NMR spectrum of LI71 Precursor (Imine) (2).

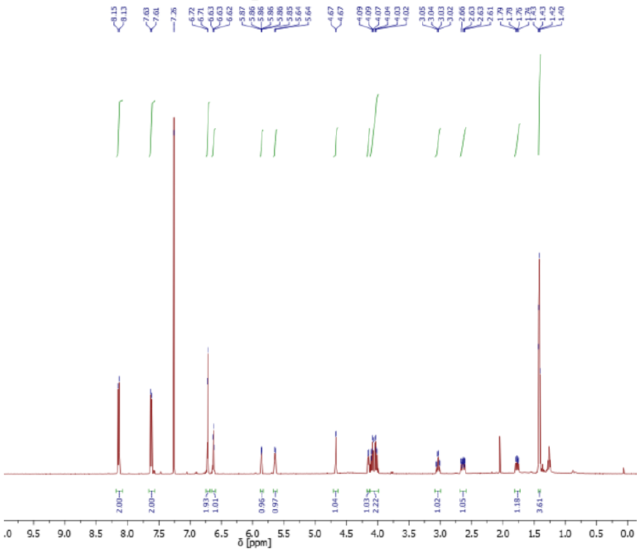


**Fig. S5: $^{1}$H NMR spectrum of LI71 (1).**

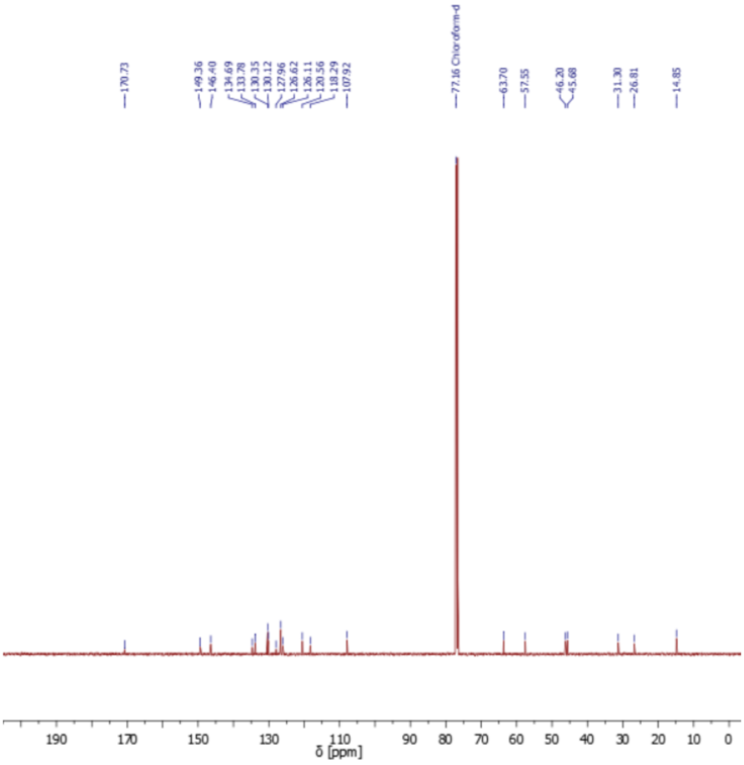


**Fig. S6: $^{13}$C NMR spectrum of LI71 (1).**

Synthetic scheme:

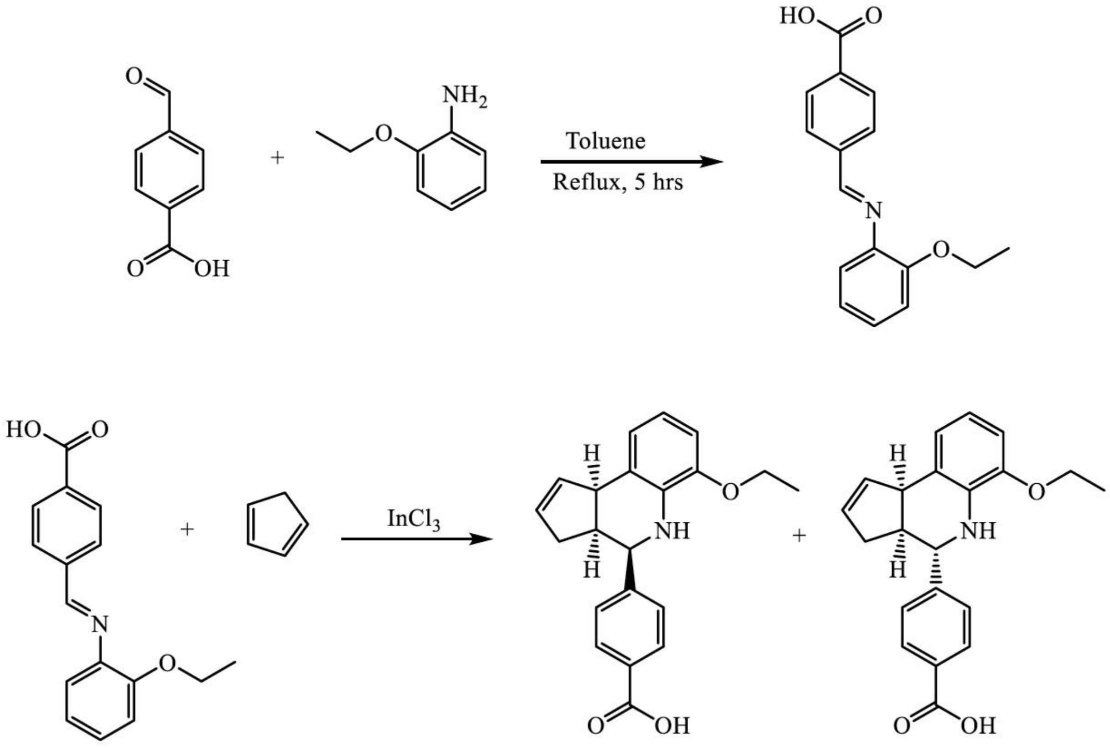


***rac*-A**

**LI71 (1)**

***rac*-B**

Step 1: Synthesis of LI71 Precursor: (E)-4-(((2-ethoxyphenyl)imino)methyl)benzoic acid

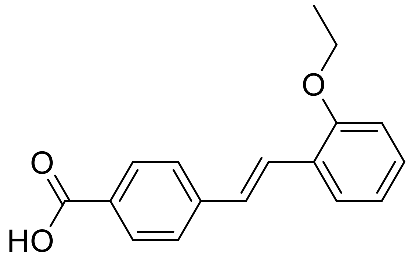

A toluene solution (10 mL) of 2-Ethoxyaniline (4.0 g, 30 mmol)) was added to a hot solution (90 mL) containing 4-Formylbenzoic acid (5.0 g, 34 mmol. The reaction mixture was heated to reflux for 5 hrs using a Dean-Stark apparatus, where upon the solution turned yellow. The product was crystallized from Acetonitrile (75 mL) giving yellow crystals, 6.0 g of pure product were obtained with a yield of 75%.

$^{1}$H NMR (400 MHz, $CDCl_3$) δ [ppm] = 10.30 (s, 1H), 8.62 (s, 1H), 8.25 (dd, J = 27.7, 8.3 Hz, 1H), 8.02 (dd, J = 13.4, 8.4 Hz, 1H), 7.23 – 7.06 (m, 1H), 6.99 (t, J = 7.0 Hz, 1H), 4.13 (dd, J = 13.8, 6.9 Hz, 1H), 1.44 (t, J = 7.0 Hz, 1H)

$^{13}C$ NMR (101 MHz, $CDCl_3$) δ [ppm] = 171.13 (O=C), 131.23 (ArC), 130.81 (ArC), 130.55 (C=C), 129.59 (C=C), 128.74 (C=C), 117.31 (d, J = 772.4 Hz), 113.37 – 107.38 (C=C), 64.58 (C-O), 14.88 (CH)

Step 2: Synthesis of LI71

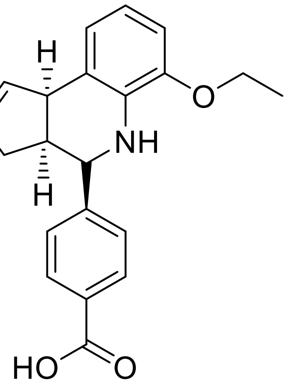


A mixture of Imine (from step 1, 3.8 g, 14 mmol), $InCl_3$ (620 mg, 2.8 mmol), and freshly cracked cyclopentadiene (2.3 ml, 28 mmol) were added to 25 mL of Acetonitrile and stirred at r.t. for 1 hr. Upon completion, the reaction mixture was washed w/ sat. $NaHCO_3$ and extracted with Dichloromethane (DCM), washed with $H_2O$, the organic layers were then dried over $Na_2SO_4$. The solvent was removed under reduced pressure, the remaining and resulting raw product (diastereomeric ratio 90/10) was then purified via column chromatography using a (70:30) Cyclohexane and Ethylacetate solvent mixture, yielding the desired product as a racemic mixture.

Product yield = 54%

$^{1}H$ NMR (500 MHz, $CDCl_3$) δ [ppm] = 8.14 (d, J = 8.3 Hz, 2H), 7.62 (d, J = 8.2 Hz, 2H), 6.72 (d, J = 5.0 Hz, 2H), 6.63 (t, J = 4.7 Hz, 1H), 5.88 – 5.83 (m, 1H), 5.64 (d, J = 4.2 Hz, 1H), 4.67 (d, J = 3.1 Hz, 1H), 4.15 (d, J = 7.5 Hz, 1H), 4.12 – 3.99 (m, 2H), 3.04 (qd, J = 9.0, 3.3 Hz, 1H), 2.68 – 2.59 (m, 1H), 1.81 – 1.72 (m, 1H), 1.41 (t, J = 6.2 Hz, 4H)

$^{13}C$ NMR (126 MHz, $CDCl_3$) δ [ppm] = 170.73 (O=C), 149.36 (ArC), 146.40 (ArC), 134.69 (C=C), 133.78 (C=C), 130.35 (C=C), 130.12 (C=C), 127.96 (C=C), 126.62 (C=C), 126.11 (C=C), 120.56 (C=C), 118.29 (C=C), 107.92 (C=C), 63.70 (C-O), 57.55 (C-C), 46.20 (C-O), 45.68 (C-N), 31.30 (C-C), 26.81 (C-C), 14.85 (C-C)

**Fig. S7:** Schematic representation for LI71 synthesis.
